# Metformin use history and genome-wide DNA methylation profile: potential molecular mechanism for aging and longevity

**DOI:** 10.1101/2022.10.22.513357

**Authors:** Pedro S. Marra, Takehiko Yamanashi, Kaitlyn J. Crutchley, Nadia E. Wahba, Zoe-Ella M. Anderson, Manisha Modukuri, Gloria Chang, Tammy Tran, Masaaki Iwata, Ryan Cho, Gen Shinozaki

## Abstract

**Background:** While there are medications that treat and manage age-related diseases, a compound that prolongs lifespan is yet to be discovered. Nonetheless, metformin, a commonly prescribed anti-diabetic medication, has repeatedly been shown to hinder aging in pre-clinical models and to be associated with lower mortality for humans, even among cancer patients. It is, however, not well understood how metformin can potentially prolong lifespan from a biological standpoint. We hypothesized that metformin’s potential mechanism of action for longevity is through its epigenetic modifications.

**Methods:** To test our hypothesis, we conducted a post-hoc analysis of available genome-wide DNA methylation (DNAm) data obtained from whole blood collected from inpatients with and without a history of metformin use. We assessed the methylation profile of 171 patients (first run) and only among 63 diabetic patients (second run) and compared the DNAm rates between metformin users and nonusers.

**Results:** Enrichment analysis from the Kyoto Encyclopedia of Genes and Genome (KEGG) showed pathways relevant to metformin’s mechanism of action, such as longevity, AMPK, and inflammatory pathways. We also identified several pathways related to delirium whose risk factor is aging. Moreover, top hits from the Gene Ontology (GO) included HIF-1α pathways. However, no individual CpG site showed genome-wide statistical significance (p<5E-08).

**Conclusion:** This study may elucidate metformin’s potential role in longevity through epigenetic modifications and other possible mechanisms of action.

## Introduction

We live in an aging society. According to the U.S. Census Bureau’s 2017 National Population Projections, 1 in every 5 residents will be in retirement age by 2030 ^1^. Subsequently, a more significant percentage of the population will endure the challenges of age-related diseases than ever before. Treatments targeting these diseases, such as dementia or cancer, at most “delay” the disease process but have a limited ability to “cure.” Therefore, there are growing interests in treating aging itself as a disease ^2^.

Considerable evidence from basic and pre-clinical models shows that several interventions, such as exercise, intermittent fasting, and even ingestion of certain compounds, can prolong lifespan. These promising compounds include rapamycin ^3,4^, resveratrol ^5-7^, NAD ^8^, and metformin ^9-11^. Our group also confirmed that inpatients using metformin had improved three-year survival rates compared to non-metformin users ^12^. In addition, our data also showed that prevalence of delirium was lower among those who were on metformin compared to those without ^12^.

The mechanism (or mechanisms) of action that rationalizes how these interventions prolong lifespan, or potentially delay aging, has been investigated heavily. Nevertheless, no exact process is well understood, especially for metformin. It is believed that epigenetics is one of the most important molecular mechanisms of aging in animals and plants; thus, it is plausible that the “life-prolonging” effects of many interventions are through modification of epigenetic processes. For example, several reports show epigenetic changes from exercise ^13^, fasting ^14^, rapamycin ^3^, resveratrol ^5^, and NAD ^8^. However, there are only a few studies investigating the direct influence of metformin on epigenetic changes ^15-17^, suggesting that information about the influence of metformin on the epigenetic profile in humans is currently limited.

To fill the gap of such knowledge, we investigated the potential influence of metformin on the epigenetic profile by testing genome-wide DNA methylation (DNAm) in whole blood samples obtained from inpatients with and without a history of metformin use.

## Results

### Demographics

173 subjects were enrolled in this study, but only 171 were included in downstream data analysis. The average patient age was 74.4 (SD= 9.8). 58 (33.9%) subjects were females while almost all the subjects were white per self-report (n= 167; 97.7%). 108 patients were non-diabetic (non-DM) while 63 were diabetic (DM). Among the DM group, 37 had diabetes with a history of metformin prescription DM(+)Met and 26 had diabetes without a history of metformin prescription DM(−)Met. Additionally, 43 (68.3%) diabetic subjects had a history of insulin use. Charlson Comorbidity Index (CCI) and body mass index (BMI) information are also included in Table 1. No variable revealed statistically significant differences between the DM(−)Met and DM(+)Met; however CCI, BMI, and insulin use were significantly higher among the DM group compared to the non-DM group, as expected.

**Table 1:**
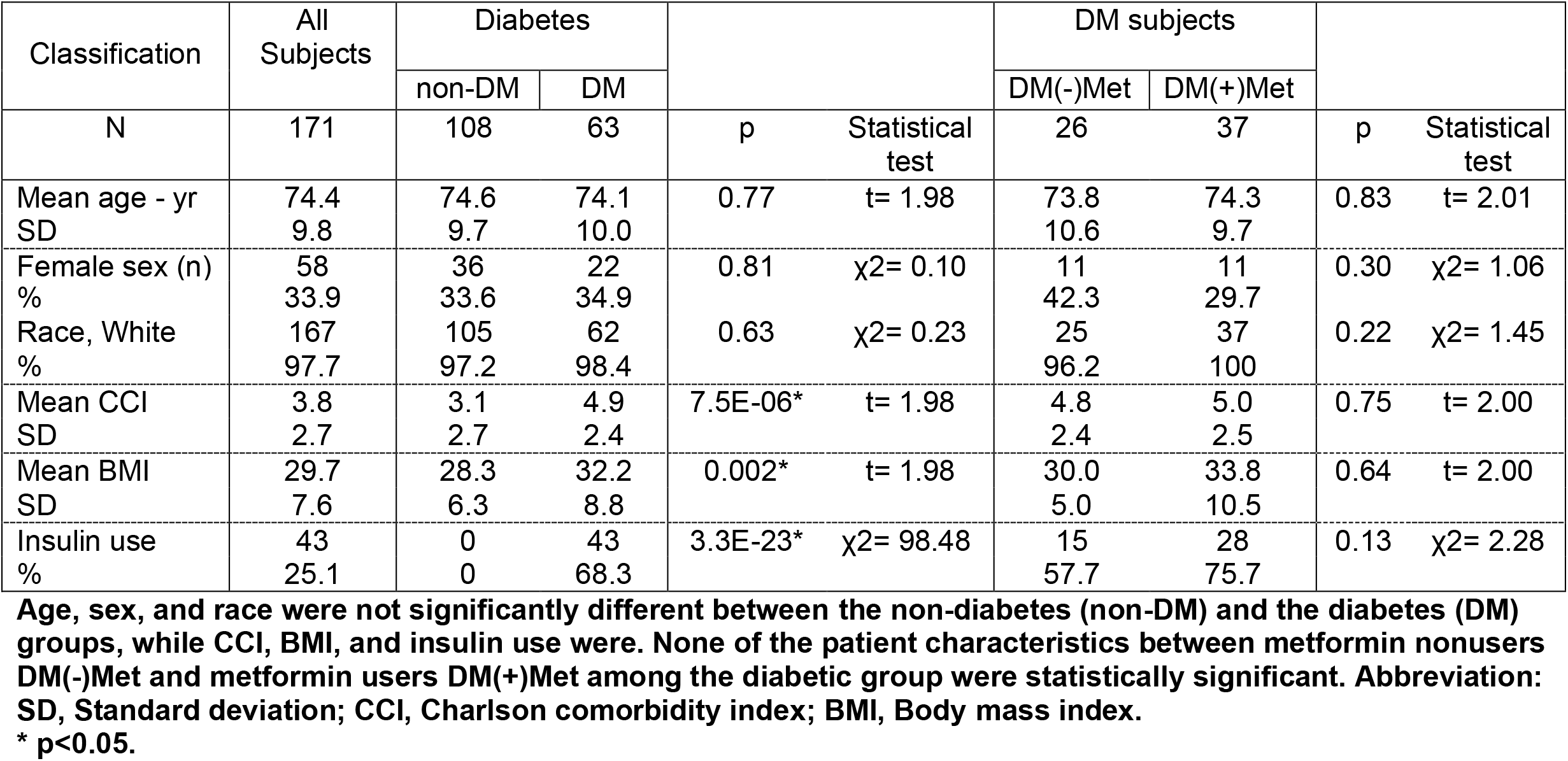
Patient characteristics.

### Met vs non-Met (including all patients regardless of diabetes status): top hits, KEGG, GO

Table 2 shows the most significant genes that differed in methylation rates between patients with and without metformin use history regardless of diabetes status (171 subjects). None of the sites met the criteria for genome-wide statistical significance (p < 5E-8).

**Table 2:**
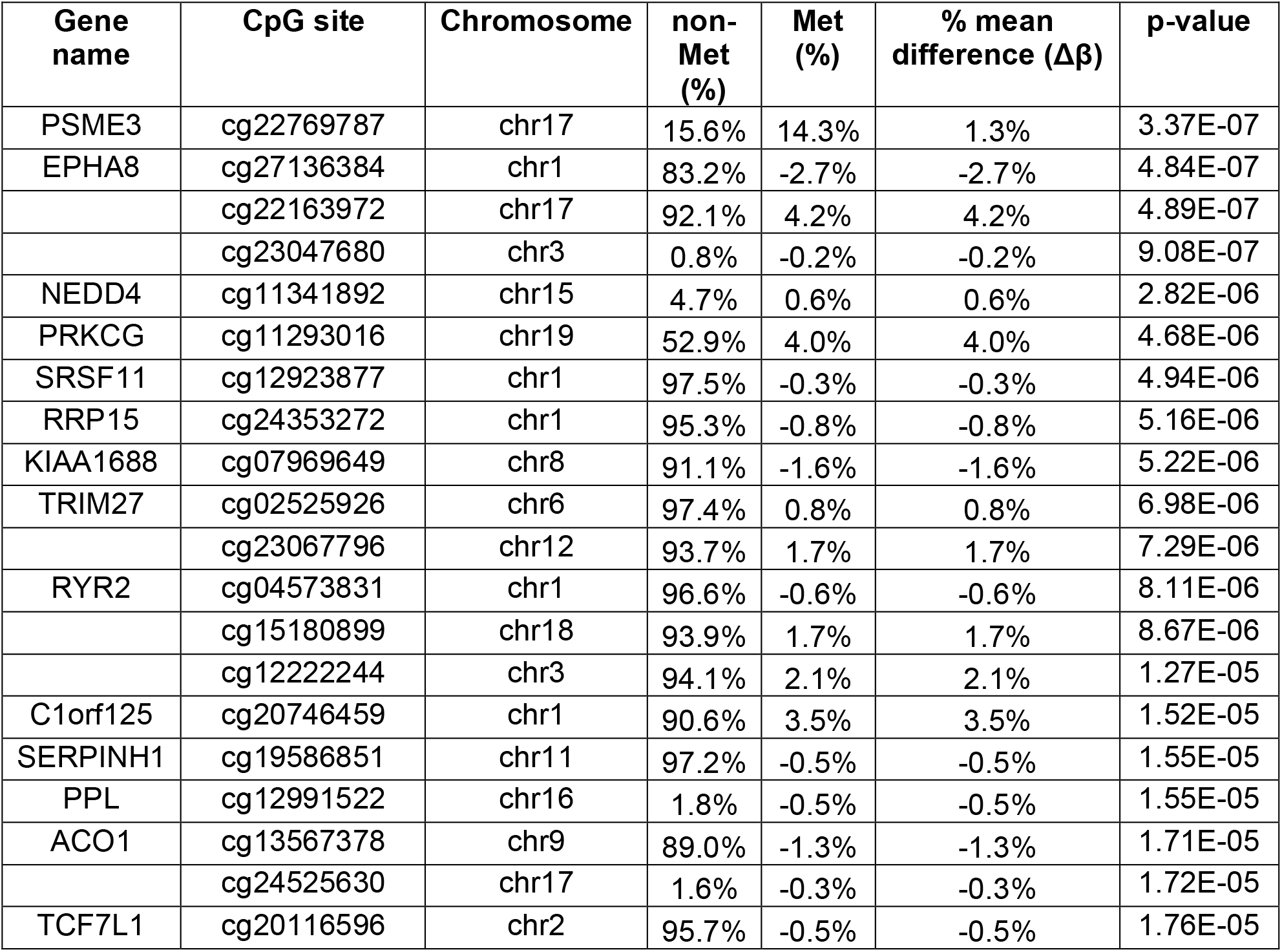
Top 20 CpG sites that differed between metformin users and nonusers among all patients.

Next, we conducted enrichment analysis using the top 330 CpG sites based on the absolute difference in methylation level (beta value) between metformin users (Met) and nonusers (non-Met) greater than 4% and the p-value less than 0.01. Enrichment analysis from the KEGG top signals showed relevant pathways to metformin’s possible roles, such as “longevity regulating pathway”, “longevity regulating pathway – multiple species”, and “AMPK signaling pathway” (Table 3). In addition, other pathways, such as “mTOR signaling pathway”, “insulin secretion”, “glutamatergic synapse”, and “circadian entrainment” were discovered (Table 3). There were also relevant pathways revealed in the GO analysis, such as “regulation of hypoxia-inducible factor-1alpha signaling pathway”, “positive regulation of hypoxia-inducible factor-1alpha signaling pathway”, and “canonical Wnt signal pathway” (Table 4), although none of the pathways in either KEGG or GO reached the False Discovery Rate (FDR) significance level (FDR<0.05) (Table 3 and 4).

**Table 3:** Top 30 KEGG pathways based on different methylation rates between metformin users and nonusers.

| **Pathway** | **N** | **DE** | **p-value** | **FDR** |
| --- | --- | --- | --- | --- |
| Relaxin signaling pathway | 129 | 6 | 0.007 | 1 |
| **Longevity regulating pathway** | 89 | 5 | 0.008 | 1 |
| **Glutamatergic synapse** | 114 | 6 | 0.008 | 1 |
| Cushing syndrome | 155 | 6 | 0.018 | 1 |
| Parathyroid hormone synthesis, secretion and action | 106 | 5 | 0.019 | 1 |
| **AMPK signaling pathway** | 119 | 5 | 0.021 | 1 |
| Signaling pathways regulating pluripotency of stem cells | 142 | 5 | 0.028 | 1 |
| Gap junction | 88 | 4 | 0.033 | 1 |
| Insulin secretion | 86 | 4 | 0.034 | 1 |
| Melanogenesis | 101 | 4 | 0.043 | 1 |
| **Longevity regulating pathway - multiple species** | 62 | 3 | 0.051 | 1 |
| Aldosterone synthesis and secretion | 98 | 4 | 0.055 | 1 |
| Chemical carcinogenesis - DNA adducts | 69 | 2 | 0.056 | 1 |
| **Circadian entrainment** | 97 | 4 | 0.058 | 1 |
| Steroid hormone biosynthesis | 61 | 2 | 0.062 | 1 |
| Thermogenesis | 219 | 5 | 0.063 | 1 |
| Bile secretion | 89 | 3 | 0.063 | 1 |
| Metabolism of xenobiotics by cytochrome P450 | 76 | 2 | 0.068 | 1 |
| Cortisol synthesis and secretion | 65 | 3 | 0.069 | 1 |
| Thyroid hormone synthesis | 75 | 3 | 0.071 | 1 |
| Wnt signaling pathway | 166 | 5 | 0.071 | 1 |
| Vasopressin-regulated water reabsorption | 44 | 2 | 0.081 | 1 |
| **Cholinergic synapse** | 113 | 4 | 0.090 | 1 |
| Retrograde endocannabinoid signaling | 141 | 4 | 0.089 | 1 |
| Estrogen signaling pathway | 137 | 4 | 0.091 | 1 |
| Mineral absorption | 60 | 2 | 0.111 | 1 |
| Gastric cancer | 149 | 4 | 0.121 | 1 |
| **mTOR signaling pathway** | 155 | 4 | 0.123 | 1 |
| Protein digestion and absorption | 102 | 3 | 0.123 | 1 |
| Ovarian steroidogenesis | 51 | 2 | 0.123 | 1 |
| Thyroid hormone synthesis | 75 | 3 | 0.071 | 1 |

Relevant pathways from KEGG <sup>59</sup> are highlighted. Abbreviations: N= number of genes included in each pathway; DE= number of Differentially Expressed genes, which are the number of genes from the top CpG sites; FDR= False Discovery Rate

**Table 4:** Top 30 GO pathways based on different methylation rates between metformin users and nonusers.

| Pathway | Ont | N | DE | p-value | FDR |
| --- | --- | --- | --- | --- | --- |
| homophilic cell adhesion via plasma membrane adhesion molecules | BP | 168 | 8 | 7.20E-04 | 1 |
| long-term synaptic depression | BP | 31 | 4 | 9.77E-04 | 1 |
| locomotory behavior | BP | 198 | 9 | 0.001 | 1 |
| midbrain dopaminergic neuron differentiation | BP | 17 | 3 | 0.002 | 1 |
| cell surface receptor signaling pathway involved in cell-cell signaling | BP | 622 | 17 | 0.002 | 1 |
| negative regulation of synaptic transmission | BP | 71 | 5 | 0.002 | 1 |
| canonical Wnt signaling pathway | BP | 335 | 11 | 0.002 | 1 |
| calcium ion binding | MF | 698 | 17 | 0.003 | 1 |
| hexose mediated signaling | BP | 6 | 2 | 0.003 | 1 |
| sugar mediated signaling pathway | BP | 6 | 2 | 0.003 | 1 |
| glucose mediated signaling pathway | BP | 6 | 2 | 0.003 | 1 |
| cellular response to acid chemical | BP | 209 | 8 | 0.003 | 1 |
| cell-cell signaling | BP | 1847 | 345 | 0.003 | 1 |
| regulation of ion transmembrane transporter activity | BP | 256 | 9 | 0.004 | 1 |
| mesoderm development | BP | 133 | 6 | 0.004 | 1 |
| neuronal cell body membrane | CC | 27 | 3 | 0.004 | 1 |
| cell body membrane | CC | 28 | 3 | 0.004 | 1 |
| regulation of transmembrane transporter activity | BP | 264 | 9 | 0.005 | 1 |
| **regulation of hypoxia-inducible factor-1alpha signaling pathway** | BP | 1 | 1 | 0.005 | 1 |
| **positive regulation of hypoxia-inducible factor-1alpha signaling pathway** | BP | 1 | 1 | 0.005 | 1 |
| cellular response to vitamin K | BP | 1 | 1 | 0.005 | 1 |
| cellular response to glucagon stimulus | BP | 25 | 3 | 0.005 | 1 |
| carbohydrate mediated signaling | BP | 8 | 2 | 0.005 | 1 |
| seminal vesicle morphogenesis | BP | 1 | 1 | 0.005 | 1 |
| glucagon-like peptide 1 receptor activity | MF | 1 | 1 | 0.005 | 1 |
| behavior | BP | 593 | 15 | 0.005 | 1 |
| nicotinamide phosphoribosyltransferase activity | MF | 1 | 1 | 0.006 | 1 |
| response to D-galactose | BP | 1 | 1 | 0.006 | 1 |
| embryonic skeletal system development | BP | 125 | 6 | 0.006 | 1 |
| regulation of transporter activity | BP | 279 | 9 | 0.006 | 1 |

Relevant pathways are highlighted. Abbreviations: Ont= Ontology; BP= biological process; CC= cellular component; MF= molecular function; N= number of genes included in each pathway; DE= number of Differentially Expressed genes, which are the number of genes from the top CpG sites; FDR= False Discovery Rate

### Met vs non-Met (including only patients with type 2 diabetes mellitus): top hits, KEGG, GO

Table 5 shows the most significant genes that differed in methylation rate between metformin users and nonusers among the diabetes group (63 subjects). Similar to the previous analysis, no gene reached genome-wide statistical significance (p < 5E-8).

**Table 5:** Top 20 CpG sites that differed between metformin users and nonusers among the diabetes group.

| Gene name | CpG site | Chromosome | non-Met (%) | Met (%) | Mean difference ($\Delta\beta$) | p-value |
| --- | --- | --- | --- | --- | --- | --- |
|  | cg19873536 | chr10 | 78.3% | 67.9% | 10.4% | 1.28E-06 |
|  | cg13596208 | chr9 | 1.9% | 2.7% | -0.9% | 2.29E-06 |
| HBA1 | cg01704105 | chr16 | 40.5% | 33.7% | 6.8% | 5.42E-06 |
| DUOX2 | cg02550961 | chr15 | 1.5% | 1.9% | -0.4% | 6.10E-06 |
| NEO1 | cg12516231 | chr15 | 2.2% | 3.2% | -0.9% | 6.97E-06 |
| C7orf46 | cg06685724 | chr7 | 2.1% | 2.9% | -0.8% | 1.28E-05 |
| NAT15 | cg00484396 | chr16 | 9.8% | 4.9% | 4.8% | 1.56E-05 |
|  | cg14685975 | chr5 | 89.9% | 92.1% | -2.2% | 1.64E-05 |
| CTSL | cg02104500 | chr9 | 3.6% | 4.9% | -1.4% | 1.66E-05 |
|  | cg12584257 | chr9 | 67.6% | 77.2% | -9.6% | 1.69E-05 |
| NAT15 | cg22508957 | chr16 | 10.9% | 6.3% | 4.6% | 1.84E-05 |
| AREL1 | cg11034672 | chr14 | 11.6% | 15.0% | -3.3% | 1.86E-05 |
|  | cg24651265 | chr10 | 1.1% | 1.7% | -0.5% | 2.12E-05 |
| CMBL | cg17467873 | chr5 | 1.7% | 2.1% | -0.4% | 2.21E-05 |
| EBF4 | cg05857996 | chr20 | 77.6% | 63.6% | 13.9% | 2.23E-05 |
|  | cg18482666 | chr2 | 95.8% | 94.8% | 1.0% | 2.39E-05 |
| HRASLS5 | cg00489394 | chr11 | 6.6% | 7.1% | -0.5% | 2.40E-05 |
| AKAP13 | cg21530087 | chr15 | 2.2% | 2.6% | -0.4% | 2.59E-05 |
|  | cg15864571 | chr3 | 93.4% | 95.0% | -1.6% | 2.67E-05 |
| FLJ35024 | cg15981195 | chr9 | 2.3% | 3.5% | -1.1% | 2.91E-05 |

The enrichment analysis was generated using consistent parameters in methylation level differences (beta>4%) and p-value (<0.01). This currently analysis, however, included 1283 CpGs. KEGG showed the many of the same signals discovered from the previous analysis, including “longevity regulating pathway”, “glutamatergic synapse”, “insulin secretion”, “circadian entrainment”, and “cholinergic synapse” (Table 6). GO also showed overlapping pathways compared to the first analysis, including “hypoxia-inducible factor-1alpha signaling pathway”, but also new pathways, such as “interleukin-8-mediated signaling pathway”, “negative regulation of leukocyte apoptotic process”, “neutrophil homeostasis”, and “neuron projection”, although these pathways did not reach the FDR significance level (FDR<0.05) (Table 7).

**Table 6:** Top 30 KEGG pathways that differed between metformin users and nonusers among the diabetes group.

| Pathway | N | DE | p-value | FDR |
| --- | --- | --- | --- | --- |
| Aldosterone synthesis and secretion | 98 | 14 | 0.001 | 0.219 |
| **Circadian entrainment** | 97 | 14 | 0.001 | 0.219 |
| Cortisol synthesis and secretion | 65 | 10 | 0.003 | 0.303 |
| Thyroid hormone synthesis | 75 | 10 | 0.004 | 0.303 |
| Regulation of lipolysis in adipocytes | 55 | 8 | 0.006 | 0.330 |
| Parathyroid hormone synthesis, secretion and action | 106 | 13 | 0.006 | 0.330 |
| Insulin secretion | 86 | 11 | 0.007 | 0.330 |
| Calcium signaling pathway | 238 | 21.5 | 0.009 | 0.388 |
| cAMP signaling pathway | 221 | 19 | 0.010 | 0.388 |
| **Cholinergic synapse** | 113 | 13 | 0.012 | 0.420 |
| Chemical carcinogenesis - receptor activation | 212 | 16 | 0.015 | 0.435 |
| **Glutamatergic synapse** | 114 | 13 | 0.016 | 0.435 |
| Rap1 signaling pathway | 210 | 19 | 0.016 | 0.435 |
| Thermogenesis | 219 | 15 | 0.020 | 0.468 |
| Amphetamine addiction | 69 | 8 | 0.021 | 0.468 |
| Neuroactive ligand-receptor interaction | 349 | 19.5 | 0.022 | 0.468 |
| Pancreatic secretion | 101 | 9 | 0.029 | 0.552 |
| Long-term potentiation | 67 | 8 | 0.029 | 0.552 |
| Cocaine addiction | 49 | 6 | 0.036 | 0.552 |
| Phospholipase D signaling pathway | 147 | 14.5 | 0.038 | 0.552 |
| cGMP-PKG signaling pathway | 166 | 14.5 | 0.039 | 0.555 |
| Apelin signaling pathway | 139 | 12 | 0.040 | 0.555 |
| Nicotine addiction | 40 | 5 | 0.040 | 0.555 |
| EGFR tyrosine kinase inhibitor resistance | 78 | 9 | 0.041 | 0.555 |
| Inflammatory mediator regulation of TRP channels | 98 | 10 | 0.042 | 0.555 |
| Gap junction | 88 | 9 | 0.042 | 0.555 |
| Type II diabetes mellitus | 46 | 6 | 0.043 | 0.558 |
| **Longevity regulating pathway** | 89 | 9 | 0.045 | 0.561 |
| Salivary secretion | 92 | 8 | 0.048 | 0.578 |
| Bladder cancer | 41 | 5 | 0.054 | 0.610 |

Relevant pathways from KEGG <sup>59</sup> are highlighted. Abbreviations: N= number of genes included in each pathway; DE= number of Differentially Expressed genes, which are the number of genes from the top CpG sites; FDR= False Discovery Rate

**Table 7:** Top 30 GO pathways that differed between metformin users and nonusers among the diabetes group.

| Pathway | Ont | N | DE | p-value | FDR |
| --- | --- | --- | --- | --- | --- |
| neuron projection | CC | 1304 | 97.1 | 1.29E-05 | 0.286 |
| second-messenger-mediated signaling | BP | 438 | 38.5 | 2.53E-05 | 0.286 |
| neutrophil homeostasis | BP | 16 | 6 | 3.77E-05 | 0.286 |
| synaptic signaling | BP | 725 | 59.5 | 9.26E-05 | 0.505 |
| trans-synaptic signaling | BP | 708 | 57.5 | 1.67E-04 | 0.505 |
| negative regulation of leukocyte apoptotic process | BP | 46 | 8 | 1.93E-04 | 0.505 |
| calcium-mediated signaling | BP | 218 | 22.5 | 1.94E-04 | 0.505 |
| chemical synaptic transmission | BP | 700 | 56.5 | 2.00E-04 | 0.505 |
| anterograde trans-synaptic signaling | BP | 700 | 56.5 | 2.00E-04 | 0.505 |
| positive regulation of cell-matrix adhesion | BP | 51 | 10 | 3.21E-04 | 0.682 |
| positive regulation of multicellular organismal process | BP | 1802 | 106 | 3.39E-04 | 0.682 |
| plasma membrane bounded cell projection | CC | 2093 | 130.1 | 4.05E-04 | 0.682 |
| interleukin-8 receptor activity | MF | 2 | 2 | 4.44E-04 | 0.682 |
| interleukin-8-mediated signaling pathway | BP | 2 | 2 | 4.44E-04 | 0.682 |
| adult behavior | BP | 144 | 17.5 | 4.73E-04 | 0.682 |
| cell junction | CC | 1858 | 123.8 | 5.16E-04 | 0.682 |
| synapse | CC | 1168 | 85.5 | 5.27E-04 | 0.682 |
| NMDA glutamate receptor activity | MF | 7 | 4 | 6.05E-04 | 0.682 |
| **hypoxia-inducible factor-1alpha signaling pathway** | BP | 6 | 3 | 6.39E-04 | 0.682 |
| regulation of dendrite development | BP | 148 | 19 | 6.44E-04 | 0.682 |
| axon | CC | 606 | 50.6 | 6.46E-04 | 0.682 |
| low voltage-gated calcium channel activity | MF | 3 | 3 | 7.18E-04 | 0.682 |
| dendrite development | BP | 232 | 26.5 | 7.28E-04 | 0.682 |
| vestibulocochlear nerve development | BP | 10 | 4 | 7.64E-04 | 0.682 |
| ionotropic glutamate receptor signaling pathway | BP | 25 | 7 | 7.69E-04 | 0.682 |
| neuron projection development | BP | 976 | 74 | 8.82E-04 | 0.682 |
| cellular response to glucose stimulus | BP | 132 | 15 | 9.26E-04 | 0.682 |
| locomotory behavior | BP | 198 | 21.5 | 9.36E-04 | 0.682 |
| cellular response to hexose stimulus | BP | 134 | 15 | 1.08E-03 | 0.682 |
| positive regulation of cellular component biogenesis | BP | 533 | 41 | 1.12E-03 | 0.682 |

**Relevant pathways are highlighted. Abbreviations: Ont= Ontology; BP= biological process; CC= cellular component; MF= molecular function; N= number of genes included in each pathway; DE= number of Differentially Expressed genes, which are the number of genes from the top CpG sites; FDR= False Discovery Rate**

### DNA methylation age acceleration

Among the diabetes group, metformin nonusers had a mean age acceleration of -8.07 compared to a mean age acceleration of -4.47 for metformin users (p = 0.11) (Figure 1). This difference was smaller among the all-subjects included regardless of diabetes status (−5.92 for metformin nonusers vs. -4.47 for metformin users; p = 0.34) (Figure 2). Both analyses did not reach statistical significance.

**Figure 1:**
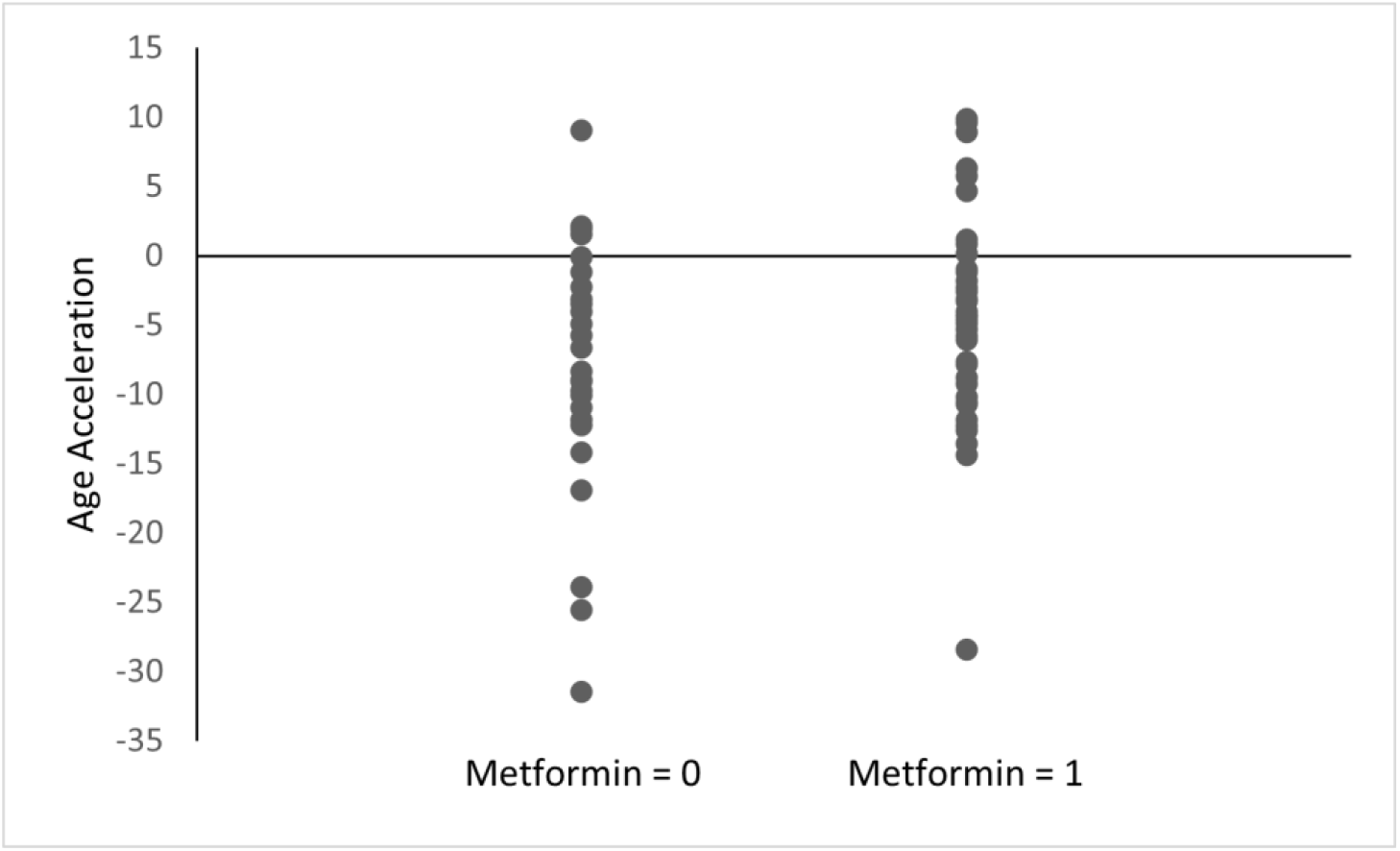
Age acceleration between metformin users and nonusers among the diabetes group. Age acceleration was calculated using the Horvath epigenetic clock as DNAm age-chronological age. Metformin = 0: without history of metformin use, Metformin = 1: with history of metformin use. p = 0.11

**Figure 2:**
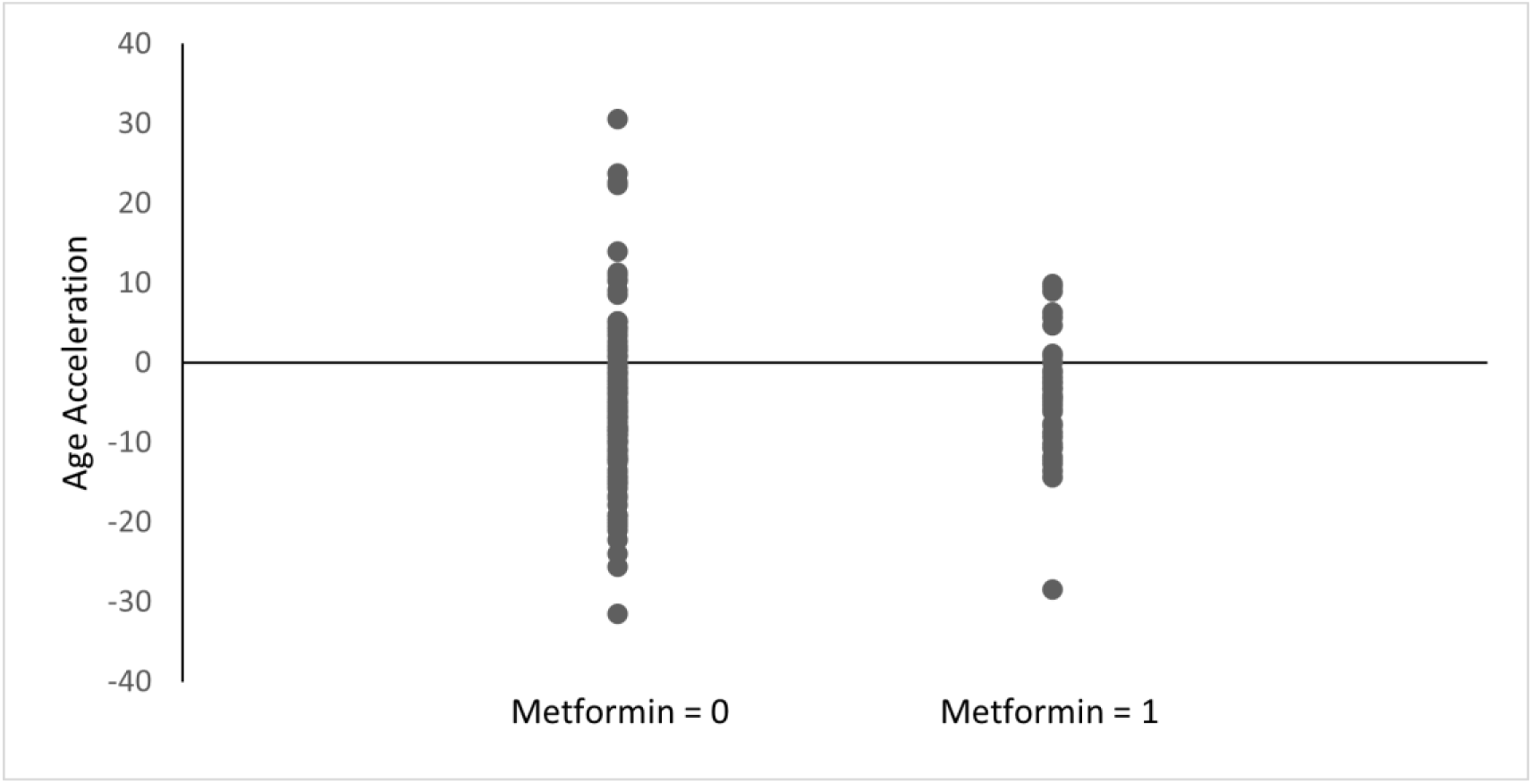
Age acceleration between metformin users and nonusers. Age acceleration was calculated using the Horvath epigenetic clock as DNAm age-chronological age. Metformin = 0: without history of metformin use, Metformin = 1: with history of metformin use. p = 0.34.

## Discussion

In this study, we compared genome-wide DNA methylation rates among metformin users and nonusers to investigate the potential epigenetic effects of metformin exposure. Enrichment analysis was employed to elucidate the possible mechanisms of action induced by metformin. Our KEGG analysis revealed evidence of differences in epigenetic profiles involved in “longevity” such as “longevity regulating pathway” and “longevity regulating pathway – multiple species” (Table 3 and 6). Although it was not statistically significant, the appearance of these pathways among top signals in the KEGG analysis demonstrates the potential role of the epigenetic processes manifesting the effect of metformin on longevity. The same KEGG analysis also showed “AMPK signaling pathway” (Table 3). AMP-activated protein kinase (AMPK), an energy sensor that regulates metabolism, is commonly referred to as one of the targets of metformin’s hypothetical mechanisms of action ^18,19^, although there is also evidence that metformin’s effects are in part AMPK-independent ^20^. Furthermore, AMPK activation is related to subsequent activation of hypoxia-inducible factors ^21^ which also appeared in our GO analyses as “regulation of hypoxia-inducible factor-1alpha signaling pathway” and “positive regulation of hypoxia-inducible factor-1alpha signaling pathway” (Table 4), as well as “hypoxia-inducible factor-1alpha signaling pathway” (Table 7). Hypoxia-inducible factor-1alpha (HIF-1α) is a transcription factor expressed in nucleated cells and mediated by oxygen levels. HIF-1α has been implicated in age-related diseases, endothelial senescence progression, AMPK, and many other pathways ^22^. Beyond metformin’s potential epigenetic medication related to longevity, several pathways related to delirium, such as “circadian entrainment”, “cholinergic synapse”, and “glutamatergic synapse”, were identified (Tables 3 and 6). These pathways are intriguing from metformin’s possible “anti-aging” standpoint as age is a major risk factor of delirium.

The beneficial effects of metformin on lifespan have been widely studied. Previous studies reported that metformin increased median lifespan of *C. elegans* co-cultured with *E*.*coli* by more than 35% ^9,23^, and prolonged the lifespan of mice ^10^. Patients with age-related diseases such as cardiovascular diseases and cancer who take metformin also had lower rates of mortality ^24,25^. Our recent study using a cohort of over 1,400 inpatients also revealed that diabetic patients with a history of metformin use have a significantly lower 3-year mortality than diabetic patients who have not taken metformin ^12^. There are, however, conflicting reports as well. For example, the same effect was not observed in *Drosophila* ^26^. Also, age-dependent, dose-dependent, and gender-dependent variable effects on lifespan were reported in mice ^27,28^. Although these previous studies’ results are not consistent, our cohort mentioned above (from which the present data are an analysis of its subgroup) clearly showed a positive influence of metformin use on survival among diabetic inpatients ^12^.

Our epigenetics data presented herein support metformin’s broad range of potential effects as indicated by the pathways identified through the enrichment analysis. The KEGG analysis (Table 7) showed several signals related to inflammation and the immune system, such as “interleukin-8 receptor activity” and “negative regulation of leukocyte apoptotic process.” The appearance of inflammation-related pathways is intriguing considering strong evidence showing that elderly people present with low-grade, chronic inflammation ^29^. These signals identified in our study may support our hypothesis that metformin can modify the inflammatory process through epigenetic modification and influence the likelihood of survival. Consistent with our data, Barath et al. also reported that metformin inhibited cytokine production from Th17 by correcting age-related changes in autophagy and mitochondrial bioenergetics, indicating its potential for the medication to promote healthy aging ^30^. Among the literature supporting metformin’s role in suppressing inflammation, clinical trials including the Diabetes Prevention Program (DPP) ^31^ and Bypass Angioplasty Revascularization Investigation 2 Diabetes (BARI 2D) ^32^ have provided further evidence of metformin’s role in changing inflammatory biomarker levels among diabetic patients, while other clinical trials, such as the Lantus for C-reactive Protein Reduction in Early Treatment of Type 2 Diabetes (LANCET) ^33^, have found opposing evidence. Although several studies mentioned here have investigated the relationship between metformin and its potential anti-inflammation, a clinical trial aimed to confirm metformin’s role in aging is yet to be seen ^34,35^. It is worth mentioning, nonetheless, a small clinical study that demonstrated the regression of epigenetic age of patients through the administration of recombinant human growth hormone (rhGH), dehydroepiandrosterone (DHEA), and metformin ^15^. As the study team administered three medications to their subjects at the same time, it is impossible to distinguish epigenetic changes caused only by metformin. It is also worth mentioning the unexpected results from the Horvath epigenetic clock since subjects with history of metformin use had relatively higher age acceleration than subjects without history of metformin. Still, neither reached statistical significance (p < 0.05). Future prospective studies comparing epigenetics marks before and after metformin use would be needed to better understand the direct effect of the medication.

In DM-only subjects, A-kinase anchoring protein 13 (*AKAP13*) gene was found (Table 5). A recent study showed that AKAP13 inhibits mammalian target of rapamycin complex 1 (mTORC1), which was present in our enrichment analysis as “mTOR signaling pathway” (Table 3). Furthermore, the degree of *AKAP13* expression in lung adenocarcinoma cell lines correlates with mTORC1 activity ^36^. Metformin’s anti-inflammatory effect has been shown to occur though eventual AMPK activation, which also inhibits the mTOR signaling pathway (*34*). Metformin’s connection to AKAP13, which has yet been fully understood, deserves further investigation.

To the best of our knowledge, our study is the largest of its kind. A smaller, previous study also investigated metformin’s effect on genome-wide DNA methylation in human peripheral blood, although their study power was limited to a sample size of 32 male subjects ^37^. Enrichment analysis in the present study revealing the longevity pathway from a hypothesis-free approach further strengthens our hypothesis that metformin exhibits its potential benefit for longevity through epigenetic processes. We also identified other relevant pathways associated with metformin’s mechanisms of action, such as the AMPK signaling pathway and HIF-1α signaling pathway ^38^.

Our study has several limitations. Although 171 subjects were analyzed retrospectively in this study, a controlled prospective study with a larger sample size would provide a better picture of the epigenetic mechanism of metformin on longevity. In addition, none of the individual CpG sites reached genome-wide significance (p < 5E-08). Thus, our findings should be interpreted as exploratory and hypothesis-generating. However, the fact that we found their biological relevance to metformin’s roles is still worth noting. As diabetes and metformin use status of the subjects was determined based on a retrospective chart review of electronic medical records, there are possibilities for misclassification, although we were still able to find multiple relevant pathways and genes of interest related to metformin’s action. Moreover, the duration of metformin use was not precisely assessed, making our definition of “metformin history use” broad since it might have included patients who took metformin for only a few months and patients who took metformin for years, for instance. Also, other types of diabetic medications were not investigated, such as sulfonylureas and glinide drugs as we used an already completed study dataset from our previous work. The rationale for us not investigating the influence of other diabetic medications was based on past literature showing that those diabetic medications other than metformin did not show benefits for survival. In fact, sometimes they were associated with worse mortality ^39-41^.

In summary, the data presented here support our hypothesis that epigenetics, especially DNA methylation, may be altered by metformin use and that such epigenetic processes potentially contribute to molecular mechanisms leading to longevity. Further careful investigation with larger sample size would be warranted.

## Methods

### Study participants and recruitment

We have previously recruited patients at the University of Iowa Hospital and Clinics (UIHC) for a separate study related to delirium from January 2016 to March 2020 ^42-45^. Among them, we used data from a subgroup of patients recruited from November 2017 to March 2020 who had blood samples collected and processed for the epigenetics analysis ^46-48^. Patients 18 years or older, who were admitted to the emergency department, orthopedics floor, general medicine floor, or intensive care unit were approached. Only those who consented, or whose legally authorized representative consented, were enlisted in the study. Written informed consent was obtained from all participants. Exclusion criteria included subjects whose goals of care were comfort measures only, those who were prisoners, or individuals with droplet/contact precautions. Further details of the study subjects and enrollment process are described previously ^42-45^.

We tested 173 subjects for genome-wide DNA methylation (DNAm) status, then conducted a post-hoc analysis of the available data to assess the influence of metformin. This study was approved by the University of Iowa Hospital and Clinics Institutional Review Board, and all procedures were compliant with the Declaration of Helsinki.

### Clinical information

Clinical variables were gathered through electronic medical chart review, patient interviews, and collateral information from family members ^42-45^. Metformin use, insulin use, and type 2 diabetes mellitus (DM) history were obtained by using the search terms “metformin”, “insulin”, and “DM” or “diabetes”, respectively ^12^. Only type 2 diabetes mellitus (DM) was included, excluding type 1 diabetes mellitus or gestational diabetes. If there was a history of metformin prescription before the study enrollment, patients were categorized as metformin users (Met). Those who were prescribed metformin after participation were not categorized as metformin users (non-Met) since the blood was obtained prior to such prescription.

### Sample collection

Blood samples were collected in EDTA tubes during patients’ hospital stay. Samples were shipped to the research laboratory and stored at -80 °C until downstream analysis as a batch.

### Sample analysis

DNA was extracted from whole blood following the MasterPure™ DNA Purification kit (Epicentre, MCD 85201). DNA passing quality control based on NanoDrop spectrometry and in sufficient amount through the Qubit dsDNA Broad Range Assay Kit (ThermoFischer Scientific, Q32850)) was selected for analysis for genome-wide DNAm status. 500 ng of genomic DNA from each sample was bisulfite-converted with the EZ DNA Methylation™ Kit (Zymo Research, D5002) and analyzed using Infinium HumanMethylationEPICBeadChip™ Kit (Illumina, WG-317-1002). The Illumina iScan platform scanned the arrays.

### Statistics and bioinformatics analysis

All analysis was conducted using R. The R packages ChAMP ^49^ and minfi ^50^ were used to process the data. Data from a total of 175 samples from 173 subjects were included for the statistical and bioinformatic analysis. DNAm levels for each CpG site were first compared between those with and without a history of metformin prescription (first run; Supplementary Table 1). Then, comparison limited among only DM patients between those with and without a history of metformin prescription was conducted to avoid potential influence of DM on DNAm status (second run; Supplementary Table 2).

During quality control processes, 2 samples from the first run and no samples from the second run were excluded based on the density analysis plots as a part of our quality control pipeline. 2 samples were also excluded because two patients had their blood collected twice. The first collected samples were included for further analysis while the second samples were excluded to maintain consistency between samples from all subjects. Therefore, 171 subjects from the first run and 63 subjects from the second run remained for the analysis. Furthermore, during the data loading process, probes were filtered out if they (i) had a detection p-value >0.01, (ii) had <3 beads in at least 5% of samples per probe, (iii) were non-CpG, SNP related, or multi-hit probes, or (iv) were located on chromosome X or Y. Beta mixture quantile dilation ^51^ was used to normalize samples, while the combat normalization method was used to correct for batch effect in the first run ^52,53^. The second run, which only included diabetic patients, was not corrected for batch effect because there were individual patients who were not part of any batches.

Top hits based on each CpG site difference were obtained through the RnBeads package using the limma method ^54,55^ and accounting for age, sex, insulin use, BMI and cell type proportions (CD8 T cells, CD4 T cells, natural killer cells, B cells, and monocytes) as covariates. DNAm Age Calculator available online ^56^ calculated the cell type proportions through the method reported previously ^57^.

After obtaining the top CpG sites, enrichment analysis followed using missMethyl ^58^ and unbalanced numbers of CpG sites on each gene were controlled using the EPIC array. Gene Ontology (GO) and Kyoto Encyclopedia of Genes and Genome (KEGG) ^59^ analysis were conducted. The number of CpG sites included in the analysis was determined by the combination of p-value and beta value cutoffs of the methylation rates of each CpG site (p<.01 and beta>0.04). Genome-wide significance was set at a p-value of less than = 5.0E-08.

The chi-square test compared the categorical data (sex, race, and insulin use) between two groups, while the Welch’s t-test compared the numerical data (age, BMI, and CCI) between two groups.

### DNA methylation aging clock analysis

To investigate whether subjects with history of metformin use had slower “age acceleration” than subjects without history of metformin use, we submitted the raw DNA methylation beta values to a publicly available tool, which includes the Horvath^56^ method. The calculated output was the difference between the DNA methylation age and the chronological age.

## List of Abbreviations

DM: diabetes mellitus
non-Met: patient without a history of metformin use
Met: patient with a history of metformin use
DM(−)Met: diabetic patients without a history of metformin use
DM(+)Met: diabetic patients with a history of metformin use.

## Declarations

## Acknowledgment

The authors thank the patients who participated in this study.

## Author Contributions

P.S.M. collected, analyzed the data and wrote the manuscript. T.Y. organized the clinical dataset and edited the manuscript. K.J.C, Z.E.M.A., M.M., G.C., and T.T. collected clinical data and biological samples, and processed them. N.E.W, K.J.C., M.I., and R.C. critically reviewed the manuscript. G.S. conceived the ideas of the study, planned its design and coordination, and wrote and edited the manuscript.

### Availability of data and materials

The datasets analyzed during the current study are available from the corresponding author on reasonable request.

### Competing interests

Gen Shinozaki is co-founder of Predelix Medical LLC and has pending patents as follows: “Non-invasive device for predicting and screening delirium”, PCT application no. PCT/US2016/064937 and US provisional patent no. 62/263,325; “Prediction of patient outcomes with a novel electroencephalography device”, US provisional patent no. 62/829,411; “Epigenetic Biomarker of Delirium Risk” in the PCT Application No. PCT/US19/51276, and in U.S. Provisional Patent No. 62/731,599. Pedro S. Marra, Takehiko Yamanashi, Kaitlyn J. Crutchley, Nadia E. Wahba, Zoe-Ella M. Anderson, Manisha Modukuri, Gloria Chang, Tammy Tran, Masaaki Iwata, Hyunkeun Ryan Cho have declared that no conflict of interest exists.

**Supplementary Table 1: Differential methylation table between metformin users and nonusers regardless of diabetes status**

Table shows top hit CpG sites in decreasing order of p-value (lower values on top), as well as the mean methylation differences (beta values). diffmeth.p.val: differential methylation p-value, mean.diff: mean difference

**Supplementary Table 2: Differential methylation table for between metformin users and nonusers among diabetes group**

Table shows top hit CpG sites in decreasing order of p-value (lower values on top), as well as the mean methylation differences (beta values). diffmeth.p.val: differential methylation p-value, mean.diff: mean difference

